# Examining the Macroecological Context of Life-History, Niche Regime, and Phylogeny in Serpentine Reptiles and Amphibians

**DOI:** 10.64898/2025.12.13.694135

**Authors:** Jared Richards

## Abstract

Life history is key to understanding macroecological patterns. Measurable characteristics are particularly critical to the study of ecology. These quantifiable traits can be compared between numerous taxonomic clades to examine how evolutionary forces have shaped biological systems. Although numerous in-depth studies have focused on assessing the life histories of mammalian and avian taxa, few scientific endeavors have addressed the diversity of measurable reproductive characteristics in other vertebrate clades. Here we present the first work examine the macroecology of life history traits in serpentine reptiles and amphibians. Both chosen clades comprise taxa that are limbless, predatory, and primarily restricted to tropical and subtropical latitudes. To test the hypothesis that phylogeny limits the extent to which life-history may overlap, the body mass, lifespan, hatchling size, and clutch size of serpentine amphibians and reptiles will be compared using statistical inference methods (t-tests). The distributions of body mass, lifespan, hatchling size, and clutch size differ among serpentine reptiles and amphibians. Linear models suggest that how hatchling and clutch size vary with body mass also differ between the two. Only the linear model examining how lifespan varies with body mass indicates that broad similarities in life histories exist between serpentine amphibians and reptiles. Our results—that little overlap exists in life-history—suggest that the same macroevolutionary processes that allow serpentine amphibian and reptile ecological niche regimes to converge are not acting on their quantifiable life-history traits.

## Introduction

Life history is key to understanding macroecological patterns. Measurable characteristics are particularly critical to the study of ecology. These quantifiable traits can be compared between numerous taxonomic clades to examine how evolutionary forces have shaped biological systems. Although numerous in-depth studies have focused on assessing the life histories of mammalian and avian taxa, few scientific endeavors have addressed the diversity of measurable reproductive characteristics in other vertebrate clades (Shine 2005). Consequently, only a handful of studies have examined the macroecology of reproductive strategy and niche regime development in ectothermic vertebrates. Of those studies, most have only evaluated well-known species within these diverse groups, such as frogs, toads, and lizards (Hanken 1999). We, therefore, have only a limited understanding of the distribution of life-history strategies employed by amphibians and reptiles—clades responsible for establishing the tetrapod body plan (Laurin 2002) and pivotal in shaping both ancient and contemporary ecosystems (Hocking et al. 2014).

In recent years, ecologists have begun to compile comprehensive datasets to collect and map basic information on the approximately 10,000 amphibian and reptile species worldwide (Hanken 1999; Uetz 2000). However, many of these data reserves have not been examined thoroughly to determine the context of species-specific trait distributions (Oliveira 2017). To address this shortcoming, I will examine the life-history traits of enigmatic and serpentine (e.g., elongated and limbless) vertebrate taxa. This includes caecilians, amphiumids, and sirens within the class Amphibia, as well as all snakes and more than a dozen genera of lizards within the class Reptilia.

Both chosen clades comprise taxa that are limbless, predatory, and largely restricted to tropical and subtropical latitudes (Gaston et al. 1995). Many of these taxa, especially those within Amphibia, are difficult to study due to cryptic lifestyles, fossorial behavior, and a preference for pristine habitats where encounters with humans are rare (Longrich et al. 2015). The fact that the behavior, morphology, and general ecology of these two diverse yet heavily understudied clades are similar—even though these groups have been distinct for many millions of years—provides an avenue for further exploration of how ecologically-imposed niche regimes and phylogeny shape the trajectory of life-history on our planet (van Tuinen and Hadly 2004).

Although these serpentine amphibian and reptile taxa have been shaped by many of the same convergent evolutionary forces, their unique histories and clade-specific characteristics likely constrain the malleability of their reproductive methods and life-history traits. For example, some groups of snakes (e.g., boas and pythons) attain considerable body mass and length, enabling them to assume dominant predatory roles in the ecosystems they inhabit (Shine et al. 1997). Since there is little evidence of any similar phenomenon occurring amongst serpentine amphibian taxa, it is expected that serpentine reptiles have, on average, higher body masses than serpentine amphibians. Lifespan, a significant determinant of the timing of reproductive events, differs between reptiles and amphibians (Allen et al., 2017). Differences in lifespan are hypothesized to exist amongst serpentine amphibians and reptiles as a result. There are also stark contrasts in the structure of oviparous amphibian and reptile eggs that likely affect life-history patterns. All reptiles (except for a few viviparous and ovoviviparous snakes such as boas and vipers) produce eggs encased in a biomineralized shell that prevents dehydration of the embryonic fluid. Amphibians, on the other hand, lay permeable eggs that must be fertilized externally. These incongruencies in ovum composition and parental behavior, alongside differences in body size (Shine 2005), likely affect hatchling size and clutch size (the number of eggs per reproductive event). To test the hypothesis that phylogeny limits the extent to which life-history may overlap, the body mass, lifespan, hatchling size, and clutch size of serpentine amphibians and reptiles will be compared using statistical inference methods (t-tests).

Studies of other taxa show that longevity, number of offspring, and offspring body size vary with body mass (Lindstedt et al., 1976; Congdon, 1987). Since relationships between these easily quantified life-history traits and body mass are well substantiated in previous studies, examining these same relationships among serpentine amphibians and reptiles may provide insight into how the life histories of serpentine taxa compare with those of other taxa. It is expected that distinct positive relationships between these life-history traits and body mass will be observed in serpentine amphibians and reptiles. Separate linear models for serpentine amphibians and reptiles will be developed to assess whether these hypothesized trends exist.

The abundance of species in the datasets used (Oliveira 2017 & Myhrvold 2015) will enable macroecological contextualization of serpentine amphibians and reptiles. This provides one of the first large-scale, high-resolution studies comparing the life-history traits of obscure taxa susceptible to recent anthropogenic activities that threaten to alter biological community structure worldwide.

## Methods

### Datasets

Amphibian life-history trait data were obtained from AmphiBIO: a worldwide species-by-trait data frame for all amphibians. This comprehensive dataset—encompassing seventeen traits for over sixty-five hundred species—was produced by compiling more than fifteen thousand literature sources (peer-reviewed papers, government documents, etc.) on the morphology, ecology, life-history, and general biology of amphibians. One hundred and ninety-three species of serpentine amphibians were included in the dataset.

Reptile life-history trait data were obtained from the Amniote Life History Database, a worldwide species-by-life-history-trait data frame for all amniotes (reptiles, mammals, birds). This dataset contains information on twenty-nine life-history parameters, compiled from over twenty-one thousand species of amniotes, based on peer-reviewed studies on their morphology, life history, and general biology. One thousand one hundred and forty-one species of serpentine reptiles were included in the dataset.

### Polyphylogeny and Data Manipulation

Serpentine amphibian and reptile taxa are polyphyletic and therefore encompass multiple taxonomic ranks. For example, all members of the amphibian order Gymnophiona (caecilians and relatives) are limbless, but only some families of the order Caudata (salamanders) are limbless. To identify all serpentine taxa within the datasets, a literature search was conducted to determine the highest taxonomic rank that could be used to correctly segregate serpentine taxa. After identification, all serpentine reptile taxa were combined into a distinct dataset. The same was done for all serpentine amphibian taxa. This removed non-serpentine taxa while segregating amphibians and reptiles for subsequent data manipulation.

Columns (i.e., species-specific data) not considered in this study were removed from each of the reduced datasets. The column names for the pertinent life-history traits (body mass, lifespan, hatchling size, and clutch size) and taxonomic levels (class, order, family, genus, and species) were standardized across the serpentine amphibian and reptile datasets. Units of measure for life-history traits were also standard for both serpentine amphibians and reptiles. All serpentine amphibians were assigned “0” in a newly created “Taxonomy” column, while serpentine reptiles were assigned “1”. This binary numerical system enabled subsequent data exploration and linear modeling. The two datasets, now congruent in structure, were combined to produce a master dataset including all 1,334 serpentine taxa considered in the study.

### Amphibian Body Mass Calculations

Only ten species of serpentine amphibians considered in the study had recorded body masses in the AmphiBIO dataset (Figure 1). However, 169 of these taxa had recorded body lengths. Previous studies have shown that body mass is strongly correlated to body length (Feldman and Meiri 2012). Body length and body size were transformed on a logarithmic scale, and a linear regression model was fitted to examine how serpentine amphibian body mass varies with body length. This linear model was used to estimate body mass for amphibian taxa whose body mass was not recorded in the AmphiBIO dataset. Since the body mass-specific data for serpentine reptiles is significantly more complete (Figure 1), estimation of body mass was only conducted on serpentine amphibians.

**Figure 1.**
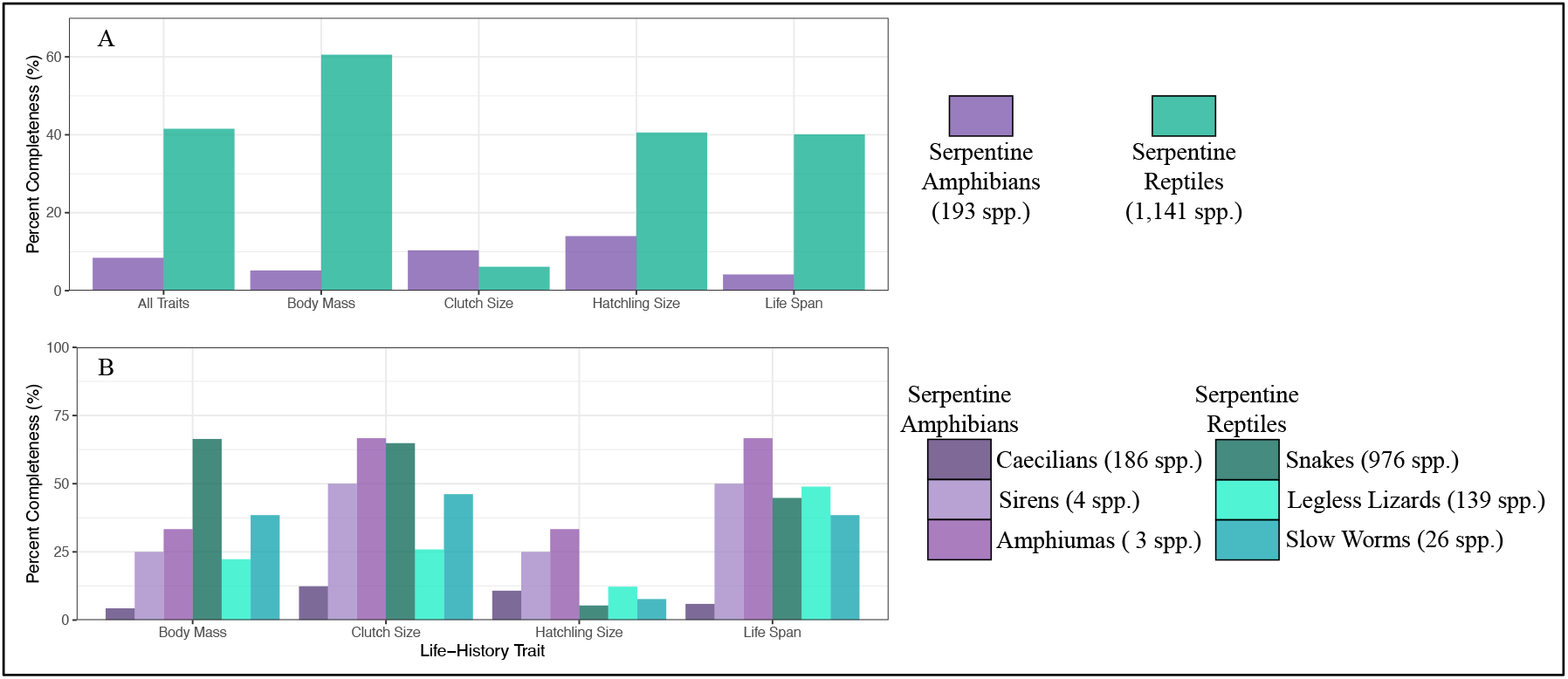
Data Completeness (%) for Serpentine Taxa (A) Data completeness (%) for serpentine amphibian and reptile life-history traits. (B) Data completeness (%) for major clades within serpentine amphibians and reptiles. The number of species in each group is included within the legend.

### Assessments of Normality

Subsequent statistical inferential testing requires an assumption of normality for data distributions. Transforming data to a logarithmic scale often reduces skewness and shifts the distribution toward normality. To determine whether the transformed or untransformed data distributions better suited the assumption of normality, Shapiro-Wilk’s tests (null hypothesis H_0_ stating that the sample came from a normally distributed population) were conducted on the untransformed and logarithmically transformed distributions for each life-history trait for both serpentine amphibians and reptiles. If the assumption of normality for a given life-history trait of either serpentine amphibians or reptiles was better substantiated by a logarithmic transformation, that life-history trait was logarithmically transformed for both groups.

### Comparing the Means of Life-History Traits

Independent two-sample t-tests were used to compare the distributions of synonymous life-history traits between serpentine amphibians and reptiles. Determining whether differences in quantifiable life-history characteristics exist between the two may allow us to make broad claims about the importance of niche regime and phylogeny in shaping life histories in these groups. Probability density plots for each trait in both serpentine groups were visualized to provide an additional avenue for examining life-history trait distributions.

### Quantifying Life-History Traits Relationships to Body Mass using Analysis of Covariance (ANCOVA)

Linear models are used to determine how lifespan, hatchling size, and clutch size vary with respect to sexually mature body mass and taxonomy (membership to the class Reptilia or Amphibia). This allows relationships among life-history traits of serpentine amphibians and reptiles to be compared. Scatter plots of the specific life-history trait versus body mass for all serpentine taxa, segregated by taxonomy, were produced to visualize and interpret the models.

## Results

### Data Completeness

91.58% of the collective species-specific data on body mass, lifespan, hatchling size, and clutch size is absent for the roughly two hundred serpentine amphibian species within the dataset (Figure 1A). Fewer than 6% of serpentine amphibians have documented lifespan and body mass data. No life-history trait is more than 14% complete in serpentine amphibians. Serpentine reptiles have been more thoroughly studied, with nearly 42% of all data on their life-history traits sampled (Figure 1A). Approximately 61% of serpentine reptiles have had their body masses documented in previous studies. More than 40% of serpentine reptiles had documented body masses, lifespans, and hatchling sizes, but only 6.13% of serpentine reptiles had documented clutch sizes (Figure 1A). When data completeness is compared amongst major clades within serpentine amphibians (i.e., caecilians, sirens, and amphiumids) and serpentine reptiles (i.e., snakes, legless lizards, and slow worms), the overall completeness of life-history traits increases (Figure 1B). Life-history data on serpentine amphibians are commonly more complete than those of serpentine reptiles when compared amongst major clades (Figure 1B).

### Amphibian Body Mass Calculations

On a log_10_-transformed scale, body length was a strong predictor of body mass for the ten serpentine amphibians (Table 1; R^2^ = 0.64) used to model the relationship. Body length and body mass for serpentine amphibians are positively correlated (P < 0.01), with body mass increasing at an exponential rate in relation to body length (Figure 2).

**Table 1.** Amphibian Body Mass Linear Model.

| Life-History Trait | $\beta_0$<br>(Intercept) | $\beta_1$<br>(Body Length) | Equation |
| --- | --- | --- | --- |
| Body Mass | -4.27* | 3.56** | $R^2 = 0.64$<br>$\text{Log}_{10}(\text{Body Mass}) = -4.27 + 3.56 * \text{Log}_{10}(\text{Body Length})$ |

The $R^2$ value indicates the proportion of variance in serpentine amphibian body mass explained by body length. Asterisks indicate the significance level of the linear model coefficients ( $P < 0.05 = *$ , $P < 0.01 = **$ , $P < 0.001 = ***$ ).

**Figure 2.**
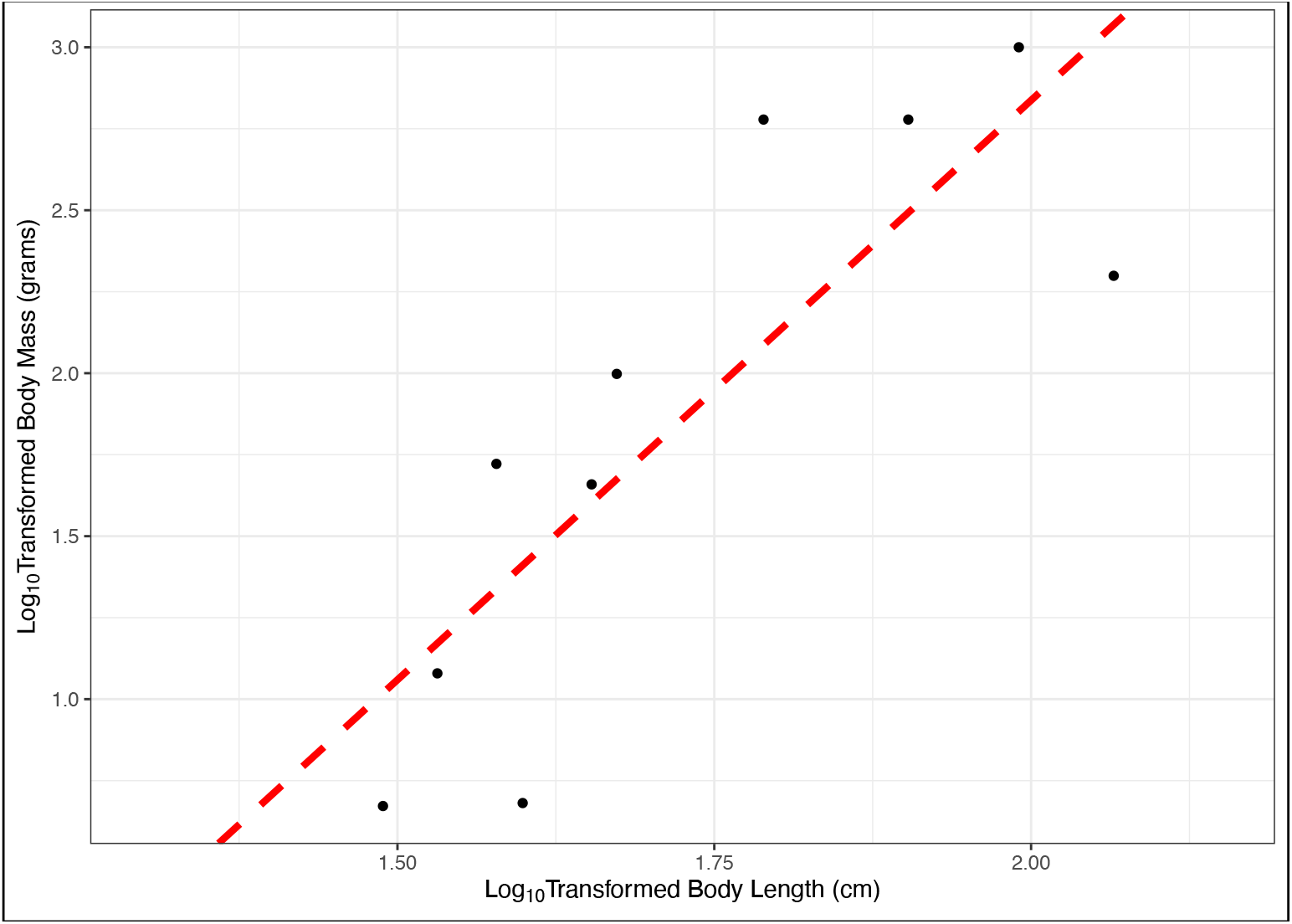
Serpentine Body Mass Estimation Visualization of the positive linear relationship between body mass and body length in the ten serpentine amphibian species with data on body mass in AmphiBIO. The given line of best fit is Log_10_(Body Mass) = -4.27 + 3.56*Log_10_(Body Length).

### Assessments of Normality

The Shapiro-Wilk tests for normality produced p-values ranging from 0.37 to less than 2.2×10^-16^, which is the smallest number greater than zero that can be stored (*e*.*g*., computational zero). For all four life-history traits of serpentine amphibians and reptiles examined in this study, logarithmic transformation increased the probability that the Shapiro-Wilk test H_0_ of normality for at least one of the two distributions was valid. Therefore, all subsequent analysis of life-history traits for both serpentine amphibians and reptiles was conducted using logarithmically transformed data. Though logarithmic transformation increased normality for most life-history trait distributions, Shapiro-Wilk tests of life-history trait distributions commonly suggested that the H_0_ of normality should be rejected (Table 2). All Shapiro-Wilk tests that provided sufficient evidence to retain the H_0_ of normality were conducted on the life-history trait distributions of serpentine amphibians. No serpentine reptile life-history trait distributions were found to suggest that the Shapiro-Wilk Test H_0_ of normality should be retained (Table 2).

**Table 2.** Life-History Trait Shapiro-Wilk Test Results.

| Serpentine Class | Life-History Trait | Shapiro-Wilk P-value for untransformed | Shapiro-Wilk P-value for $\log_{10}$ transformation | Does transformation increase normality? |
| --- | --- | --- | --- | --- |
| Amphibia | Body Mass | $5.7 \times 10^{-3***}$ | 0.37 | Yes |
| Reptilia | Body Mass | $<2.2 \times 10^{-16***}$ | $8.4 \times 10^{-7***}$ | Yes |
| Amphibia | Lifespan | 0.16 | 0.22 | Yes |
| Reptilia | Lifespan | $3.0 \times 10^{-9***}$ | $<2.2 \times 10^{-16***}$ | No |
| Amphibia | Hatchling Size | $9.1 \times 10^{-4***}$ | 0.35 | Yes |
| Reptilia | Hatchling Size | $4.4 \times 10^{-5***}$ | 0.01* | Yes |
| Amphibia | Clutch Size | $4.3 \times 10^{-9***}$ | 0.02* | Yes |
| Reptilia | Clutch Size | $<2.2 \times 10^{-16***}$ | $1.0 \times 10^{-7***}$ | Yes |

P-values for Shapiro-Wilk tests of normality are included here for all life-history trait distributions examined. Asterisks delineate the significance level of the test ( $P < 0.05 = *$ ; $P < 0.01 = **$ ; $P < 0.001 = ***$ ). A comparison of untransformed and $\log_{10}$ -transformed p-values was used to determine whether transformation brought the distributions closer to normality.

### Comparing the Means of Life-History Traits

All life-history trait distributions are significantly different for serpentine amphibian and reptile taxa (Table 3). The average mass of serpentine reptiles was higher than the average mass of serpentine amphibians (Table 3). The distribution of serpentine reptile mass is unimodal (Figure 1A); reptile mass ranges from less than 10 grams to more than 100 kilograms. The distribution of serpentine amphibian mass is unimodal. Serpentine amphibian mass ranges from hundreds of milligrams to ten kilograms (figure 3A). The average lifespan for amphibians is higher than that of reptiles (Table 3). The distribution of log_10_-transformed reptile lifespan is heavily skewed towards smaller values, and serpentine reptile lifespan ranges from less than a year to roughly one hundred years (Figure 3B). The range for log_10_-transformed serpentine amphibian lifespan is bimodal and is less skewed than that of serpentine reptiles (Figure 3B).

**Table 3.** Life-History Trait Distribution and Comparative Student T-Test Results.

| Life-History Trait (log<sub>10</sub> scale) | Amphibian Average | Amphibian Mode(s) | Reptile Average | Reptile Mode(s) | Student t-test P-value |
| --- | --- | --- | --- | --- | --- |
| Body Mass (grams) | 1.27 | 0.8 | 2.52 | 2.9 | 6.4x10<sup>-3***</sup> |
| Lifespan (years) | 1.17 | 0.75, 1.35 | 1.09 | 1.1 | 1.4x10<sup>-6***</sup> |
| Hatchling Size (cm) | -0.15 | 0.9 | 1.21 | 0.6, 1.30 | < 2.2x10<sup>-16***</sup> |
| Clutch Size | 1.30 | 1.25 | 0.84 | 0.9 | 6.4x10<sup>-5***</sup> |

Distribution-specific data included for all life-history traits of serpentine amphibians and reptiles. Student's t-test comparing logarithmically transformed synonymous life-history characteristics. Asterisks delineate the significance level of the test (P < 0.05 = \*; P < 0.01 = \*\*; P < 0.001 = \*\*\*).

**Figure 3.**
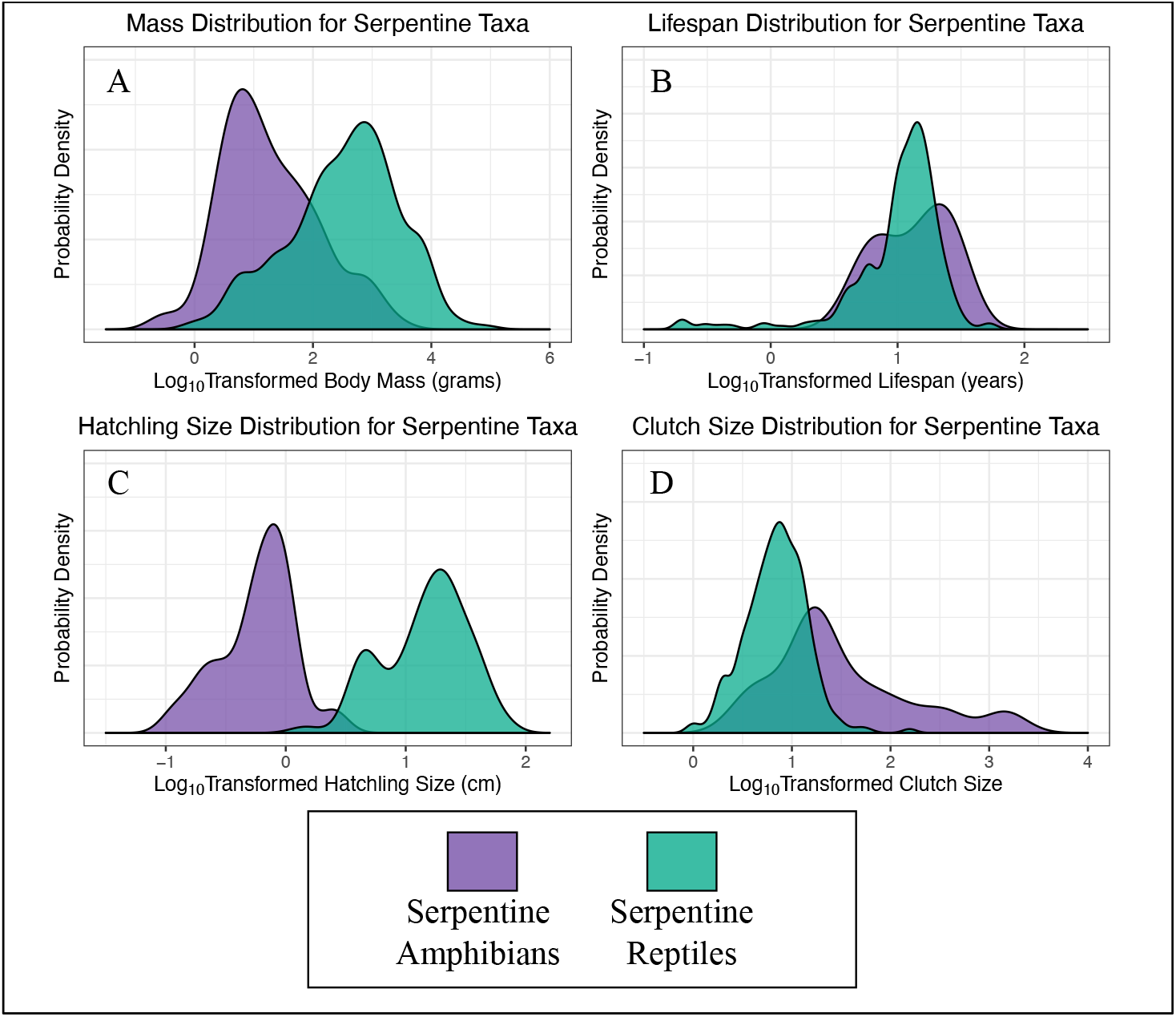
Life-History Trait Distributions Visualization of the distribution of life-history traits for serpentine amphibians and reptiles. (A) Body mass distributions; (B) Lifespan distributions; (C) Hatchling size distributions; (D) Clutch Size distribution.

Serpentine amphibian lifespans range from approximately 10 to 20 years. The average hatchling size for serpentine reptiles is larger than the hatchling size of serpentine amphibians (Table 3). The distribution of serpentine amphibian hatchling size is unimodal and ranges from millimeters to almost ten centimeters (Figure 3C). The distribution of serpentine reptile hatchling size is bimodal. Serpentine reptile hatchling size ranges from a few centimeters to nearly a meter (figure 3C). The average clutch size for serpentine amphibians is larger than the average clutch size of serpentine reptiles (Table 3). The distribution of serpentine amphibian clutch size is unimodal and skewed towards larger values (Figure 3D). Serpentine amphibian clutch size ranges from less than ten eggs to more than a thousand eggs per clutch. The distribution of serpentine reptile clutch size is more normally distributed than that of serpentine amphibians and ranges from less than ten eggs to tens of eggs per clutch (Figure 3D).

### Quantifying Life-History Traits Relationships to Body Mass using ANCOVA

Lifespan is not readily predicted by body mass and taxonomy (Figure 4A, R^2^ = 0.07). There is a weak negative relationship between lifespan and body mass for serpentine amphibians (Table 4). There is a positive relationship between lifespan and body mass for serpentine reptiles that is not significantly different than that of serpentine amphibians (table 4, P > 0.05). The intercept for serpentine reptile lifespan is lower in magnitude than that of serpentine amphibians, but not statistically distinct (Figure 4A, P > 0.05). Hatchling size is readily predicted by body mass and taxonomy (R^2^ = 0.89, Figure 4B). There is a strong positive relationship between hatchling size and body mass for serpentine amphibians (Table 4, P < 0.05). There is a positive relationship between lifespan and body mass for serpentine reptiles that is not significantly different than that of serpentine amphibians (table 4, P > 0.05). The intercept for serpentine reptile hatchling size is higher than that of serpentine amphibians and significantly different (Table 5, Figure 4B, P < 0.05). Clutch size is predicted by body mass and taxonomy (Figure 4C, R^2^ = 0.46). There is a positive relationship between clutch size and body mass for serpentine amphibians (Table 5). There is a positive relationship between clutch size and body mass for serpentine reptiles that is distinct from that of serpentine amphibians (Table 4, P < 0.05). The intercept for serpentine amphibian clutch size is lower than that of serpentine amphibians and significantly different (Figure 4C, P < 0.05).

**Table 4.**
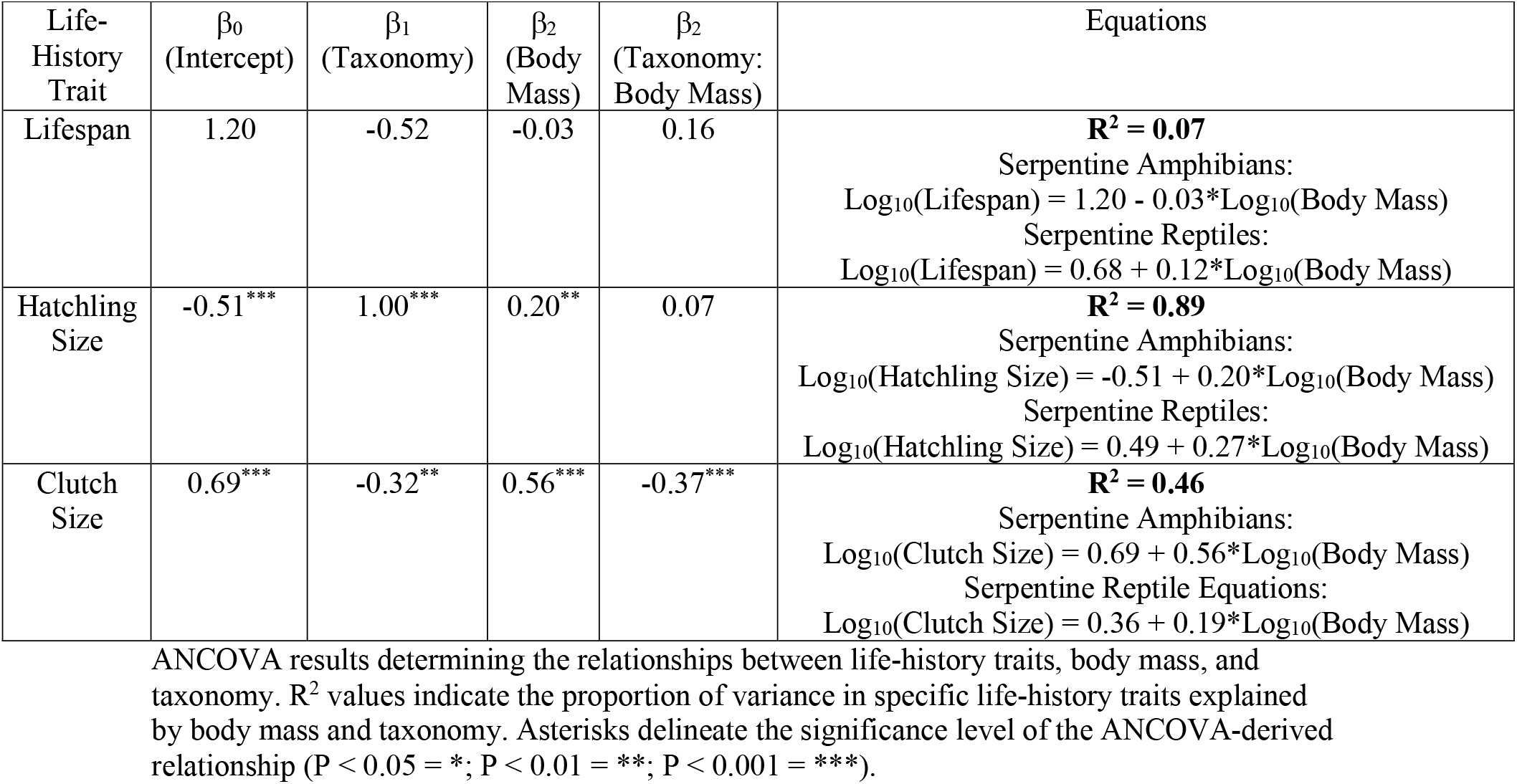
Body Mass Dependent Life-History ANCOVA Models.

**Figure 4.**
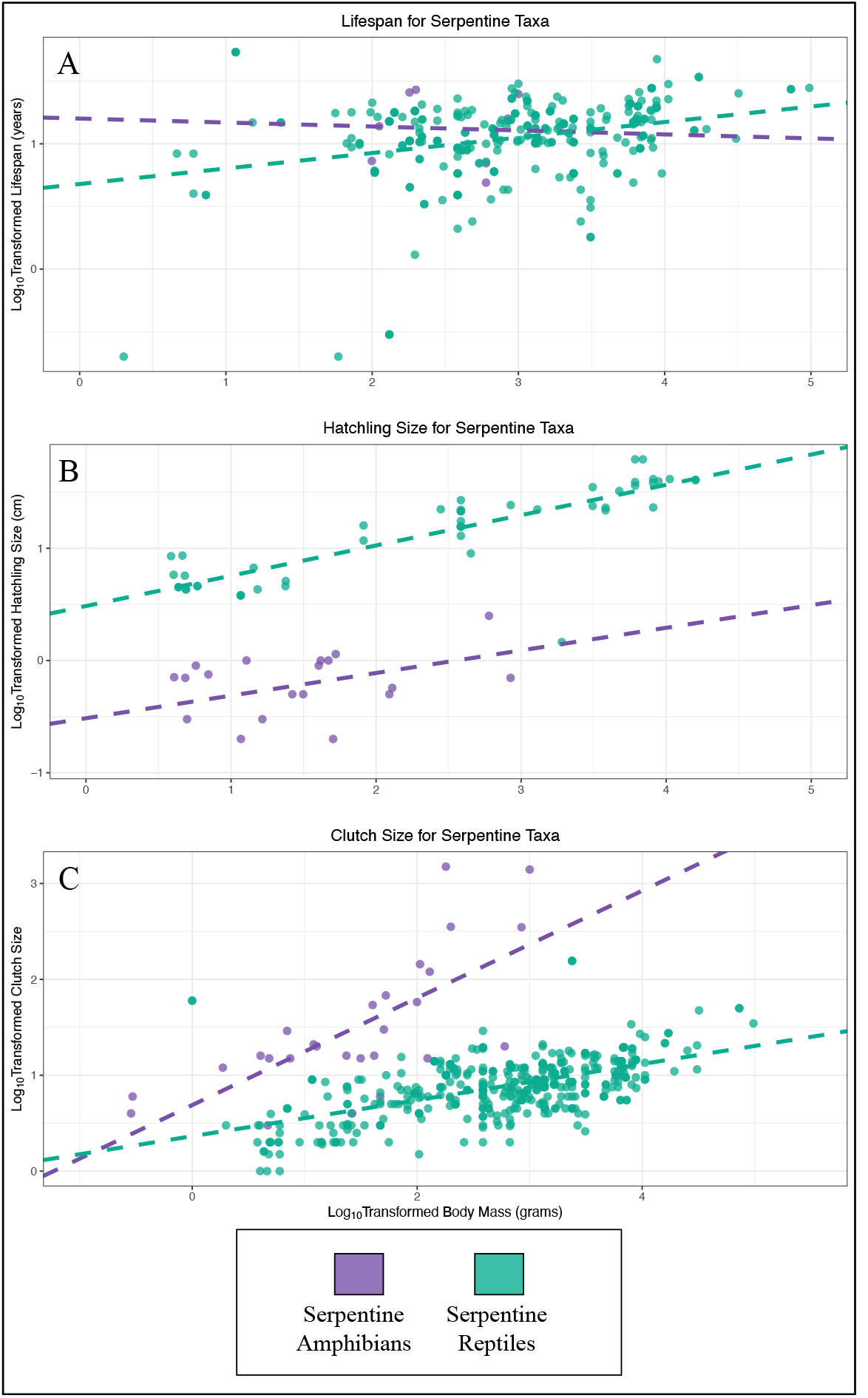
Life-History Trait Relationships to Body Mass Visualization of ANCOVA results for serpentine amphibian and reptile life-history traits. (A) Lifespan and Body Mass ANCOVA; (B) Hatchling Size and Body Mass ANCOVA; (C) Clutch Size and Body Mass ANCOVA.

## Discussion

### Data Completeness and Effect on Data Interpretation

The low percentage of data completeness observed among focal taxa, especially amphibians, corroborates claims that we know very little about the general biology and life histories of cryptic serpentine vertebrate taxa worldwide. Within-clade-specific data appear more complete than general data, mainly because some groups encompassed very few species. For example, though amphiuma-specific lifespan data is 67% complete, there were only three species of amphiuma included in the AmphiBIO dataset. Although the completeness of life-history data across major clades of serpentine amphibians and reptiles may appear comparable, sirens and amphiumas provide significantly fewer data points than more populous and well-studied clades, such as snakes and legless lizards.

The lack of serpentine amphibian-specific data may also play an essential role in interpreting the results. For example, an *a posteriori* examination of the composition of the ten serpentine amphibian species used to model the relationship between body length and body mass revealed that eight were caecilians and two were salamanders (one siren and one amphiuma). This small sample size does not fully represent the breadth of serpentine amphibian taxa and likely introduces systematic bias into the results. Caecilians and serpentine salamanders may have slightly different allometric relationships between body length and body mass that were not duly represented because the lack of data that exists (Sorenson 2004). The strong skew toward caecilians may have produced a regression that inaccurately modeled the body mass-body length relationships of serpentine salamanders, as in caecilians. Although potentially biased, the highly significant relationship observed between serpentine amphibian body mass and body length provides essential insight into the growth patterns of these taxa. The same strong allometric trends observed at a macroecological scale among other taxa (Feldman and Mieri 2012) are clearly present among poorly studied, cryptic taxa that occupy specific ecological niches.

### Life-History Trait Relationships

Most life-history trait distributions were brought closer to normality by a logarithmic transformation. The fact that a transformation did not normalize the serpentine reptilian lifespan suggests that a wide range of lifespans is present within this group. Although a large portion of serpentine reptiles have lifespans of approximately 10 years, the distribution is skewed toward shorter lifespans by a variety of species that have lifespans of only a few years. This strong skew, not seen in any other life-history trait examined in the study, suggests that a wide breadth of distinct lifespans—possibly a result of numerous phylogenetic and ecological factors—exist amongst serpentine reptiles.

Given that some snakes, such as boas and pythons, can weigh more than two hundred kilograms (Feldman et al. 2015), it was anticipated that serpentine reptiles would be almost eighteen times heavier than serpentine amphibians on average. Although the locations of body mass data differ between the two, the distributions are very similar in shape. This provides evidence that, although phylogeny and evolutionary history constrain each group, similar proportions of taxa within each group can successfully diverge from the most common body masses. Serpentine amphibians, which are generally smaller than serpentine reptiles, have longer lifespans. Additionally, lifespan declines with body mass in serpentine amphibians, contradicting numerous previous studies indicating a strong positive correlation between body mass and lifespan across the animal kingdom (Trites and Pauly 1998; Lindstedt et al. 1976). Our findings may be attributable to an inadequate sample size. If life histories were better documented, the well-substantiated positive correlation between lifespan and body mass would be observed among serpentine amphibians as well. Lifespans of roughly six years and twenty-two years are extremely common amongst serpentine amphibians. This is either a product of poor sample size or suggests that these two lifespan strategies provide ideal temporal windows for procreative events amongst amphibians.

As expected, serpentine reptiles were found to be larger than serpentine amphibians upon hatching. This aligns with previous studies showing that larger organisms produce larger offspring (Congdon and Gibbons 1985; Fischer et al. 2002) and that the shell-bound amniotic eggs of reptiles provide an excellent environment for developing larger, increasing mobility, and decreasing predation risk (Shine 2005). Though serpentine reptiles and amphibian hatchling size differ at any given body mass, there is no evidence that the positive relationship between body mass and hatchling size is not synonymous for both. This suggests that the allometry of hatchling size has been constrained by similar forces for both serpentine amphibians and reptiles. Lifespans of roughly four years and twenty years are extremely common amongst serpentine reptiles. Since this trait is well documented in serpentine reptiles, it suggests that these two lifespan strategies provide optimal temporal windows for major reproductive events. As amphibians are known to mass spawn—sometimes producing thousands of eggs at a time (Allen et al. 2017)—the fact that serpentine amphibian clutch sizes were almost three times as large as those of serpentine reptiles conforms to previously established patterns of life-history for the two groups. Serpentine amphibians of a specific mass will produce more viable offspring in any one reproductive event when compared to a serpentine reptile of synonymous body mass. The positive relationship between serpentine amphibian clutch size and body mass is greater (i.e., increases faster) than that of serpentine reptiles. This contrast in clutch size, alongside patterns observed in hatchling size, suggests that the two groups may be investing differentially in the quantity and quality of offspring.

### Niche Regime, Phylogeny, and Future Directions

The distributions of body mass, lifespan, hatchling size, and clutch size differ among serpentine reptiles and amphibians. Linear models suggest that how hatchling and clutch size vary with body mass also differ between the two. Only the linear model examining how lifespan varies with body mass indicates that broad similarities in life histories exist between serpentine amphibians and reptiles. However, drawing any major claims from this is ill-advised, as there is very little documentation of serpentine amphibian lifespan. Our results—that little overlap exists in life-history—suggest that the same macroevolutionary processes that allow serpentine amphibian and reptile niche regimes to converge are not acting on their quantifiable life-history traits. Studies of other taxa show a similar pattern. Although distinct phylogenies have converged anatomically and behaviorally under ecological and evolutionary pressures, their unique histories constrain how similar their reproductive methods may become (Emory and Clayton 2004; Chen et al. 1997).

In conclusion, we have found evidence of large-scale differences in quantifiable life histories among vertebrate taxa that are commonly excluded from present-day macroecological studies. Future studies of serpentine amphibians and reptiles should focus on filling gaps in our knowledge of their life histories and general biology. Understanding the macroecological relevance of serpentine vertebrates may enable scientists to identify the measures needed to protect these cryptic and misunderstood groups from anthropogenic disturbance.

